# Modulating Protein Function through Genetically Encoded Oxidative Chemistry

**DOI:** 10.1101/2025.11.06.686050

**Authors:** Hengyu Li, Alen Pavlič, Noor E. Ibrahim, Di Wu, Mikhail G. Shapiro

## Abstract

Oxidative chemistry underlies many endogenous signaling pathways but remains underutilized as a programmable strategy for regulating protein function in living cells. Here we establish genetically encoded oxidative chemistry as a tunable framework for modulating diverse proteins by coupling a photosensitizer to defined intracellular contexts. Using miniSOG to generate reactive oxygen species (ROS), we show that controlled intracellular oxidation increases the fluorescence of the redox reporter HyPerRed and activates redox-sensitive TRP ion channels, with strong responses in TRPA1 and TRPV1 but not TRPV4. Pathway-selective scavengers reveal differential coupling of soluble and membrane targets to distinct oxidative processes, supporting selectivity by context rather than uniform oxidative perturbation. Modulation strength and kinetics are quantitatively tunable through illumination parameters, expression ratios, and subcellular localization, with membrane targeting enhancing coupling to membrane effectors. Finally, fusion targeting of miniSOG to TRPV1 and modulation of endogenous TRPA1 in human fibroblasts extend this approach to protein-proximal and native cellular settings. Together, these results position genetically encoded oxidative chemistry as a versatile and spatially organized modality for engineering protein function in living cells within a defined operating regime.

## Introduction

Reactive oxygen species (ROS) regulate numerous cellular signaling pathways through oxidative chemistry, including cysteine and methionine oxidation^1,2^, metal-center redox regulation^3^, and lipid oxidation^4^. Despite this central role in biological signaling, oxidative chemistry remains relatively underexplored as a programmable design principle for regulating protein function in living cells. Importantly, ROS comprise chemically distinct species with different reactivities and diffusion behaviors, suggesting that effective deployment of oxidative chemistry as a regulatory strategy requires consideration of both chemical identity and spatial context.

Genetically encoded photosensitizers such as miniSOG, KillerRed, and related variants provide versatile means to trigger oxidative chemistry inside living cells^5–10^. These proteins have been widely used for chromophore-assisted light inactivation (CALI)^6,8–11^, imaging^12–17^, and proximity labeling^18–22^, where ROS are primarily exploited to damage, crosslink, or tag nearby biomolecules. However, these applications largely harness the destructive or labeling capabilities of ROS rather than their potential as a programmable chemical mechanism for regulating protein function through quantitatively controllable and spatially tunable oxidative modulation. Repurposing oxidative chemistry as a programmable regulatory modality could enable new strategies for engineering protein function in living cells. Moreover, because ROS encompass multiple chemically distinct species, how different oxidative pathways selectively engage specific protein targets remains insufficiently understood.

Here, we show that genetically encoded ROS generation can modulate protein function through oxidative chemistry in living cells and provide a strategy for implementing oxidative modulation with genetically encoded photosensitizers. We find that intracellular ROS generated by miniSOG increase HyPerRed^23^ fluorescence and activate TRP ion channels, with strong responses in TRPV1 and TRPA1 but not TRPV4. Pathway-selective scavengers further reveal differential coupling of soluble and membrane targets to distinct oxidative pathways.

Because ROS are inherently reactive and diffusible, the spatial relationship between oxidant source and target protein represents a critical design parameter for oxidative modulation. Consistent with this principle, membrane targeting of miniSOG enhances functional coupling to membrane effectors, and fusion-based targeting of miniSOG to TRPV1 enables protein-proximal ROS generation at the channel. We additionally demonstrate modulation of an endogenously expressed ion channel in its native cellular context. Together, these results demonstrate genetically encoded oxidative chemistry as a framework for engineering quantitatively tunable and spatially organized modulation of protein function in living cells within a defined operating regime.

## Results

### miniSOG-derived ROS modulate HyPerRed fluorescence in living cells

To test whether genetically encoded ROS generation could be used to modulate protein function, we first examined a soluble redox reporter. We co-expressed the photosensitizer miniSOG and the fluorescent redox reporter HyPerRed in HEK293T cells by transient co-transfection of the two plasmids (**Figure 1A**). Mock control cells were co-transfected with HyPerRed and the empty vector used for the miniSOG construct. During imaging, the culture dish was maintained in a 37°C water tank to minimize temperature fluctuations during photo-stimulation while recording HyPerRed fluorescence in real time. We chose HyPerRed because its excitation and emission spectra do not overlap with those of miniSOG, minimizing optical crosstalk during illumination and imaging^5,23^, and because it contains a reactive cysteine pair that undergoes oxidation-dependent fluorescence changes in response to local redox conditions. We therefore hypothesized that ROS generated by miniSOG upon blue-light illumination would perturb this redox equilibrium and alter HyPerRed fluorescence.

**Figure 1.**
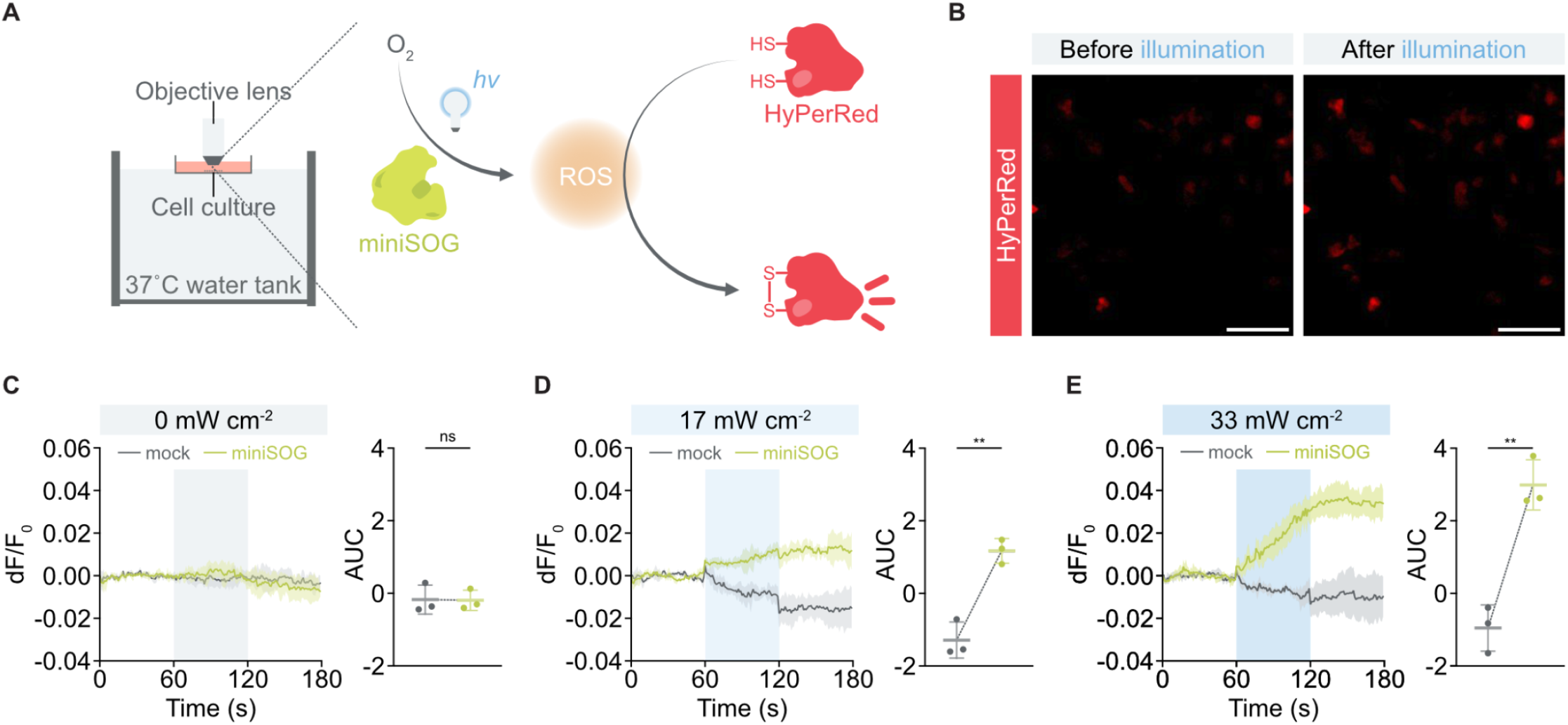
miniSOG-derived ROS modulate HyPerRed fluorescence in living cells. **(A)** Schematic of the experimental setup. HEK293T cells co-expressing miniSOG and HyPerRed were placed in a 37°C water bath during imaging. Blue-light illumination (470 nm) of miniSOG generated ROS that altered HyPerRed fluorescence. **(B)** Representative fluorescence images of HyPerRed-expressing cells before and after blue-light stimulation (33 mW cm^−2^, 60 s). Scale bar, 100 μm. **(C-E)** Time-course traces (left) and corresponding AUC quantification (right) of HyPerRed fluorescence in mock control cells (HyPerRed + empty vector) and miniSOG-expressing cells. Blue shading indicates illumination period (60-120 s). No significant change was observed at 0 mW cm^−2^ **(C)**, whereas graded increases were observed at 17 mW cm^−2^ **(D)** and 33 mW cm^−2^ **(E)**. Data are mean ± s.d. (n = 3 independent imaging dishes). Statistical significance was determined by two-tailed *t* tests; ns, not significant; **, *p* < 0.01.

Blue-light stimulation during a 60-second exposure window induced a pronounced increase in HyPerRed fluorescence in miniSOG-expressing cells, but not in mock controls (**Figure 1B-E**). Single-cell fluorescence traces were extracted from segmented HyPerRed-positive cells and averaged within each imaging dish to generate a dish-level trace representing each biological replicate. The magnitude of the response scaled with illumination intensity, exhibiting graded increases at 17 and 33 mW cm^−2^ but not at 0 mW cm^−2^ (**Figure 1C-E**). Quantification of the fluorescence area under the curve (AUC) confirmed a significant enhancement in miniSOG-expressing cells relative to mock controls. Statistical analysis further showed that fluorescence responses in miniSOG-expressing cells depended on light intensity, whereas those in controls did not (**Supplementary Figure 1**). Because responses were quantified as normalized ΔF/F_0_ values, variability in basal HyPerRed fluorescence across cells did not affect the measured responses. After illumination ended, fluorescence reached a steady level that persisted throughout the recording period. Together, these results show that miniSOG can generate a tunable intracellular oxidative input that is sufficient to modulate the fluorescence of a soluble redox-sensitive protein in living cells, providing a starting point for testing oxidative modulation in more complex protein targets.

### miniSOG-derived ROS activate redox-sensitive ion channels TRPV1 and TRPA1 in living cells

Having shown that miniSOG-generated ROS can modulate a soluble redox-sensitive reporter, we next asked whether this approach could also regulate membrane ion channels that transduce oxidative cues into ionic signals. We chose TRPV1, TRPV4, and TRPA1 because they have all been reported to respond to oxidative stress, but differ in the extent and mechanism of their redox sensitivity^24–31^, providing a physiologically relevant test set for whether miniSOG-generated ROS gate membrane proteins differentially in living cells.

We transiently co-expressed miniSOG with TRPV1, TRPV4, or TRPA1 together with the Ca^2+^ indicator jRGECO1a^32^ in HEK293T cells (**Figure 2A**). Blue-light illumination of miniSOG was used to generate ROS, and jRGECO1a fluorescence was recorded in real time as a readout of channel-mediated Ca^2+^ influx. This configuration enabled us to quantify channel-dependent Ca^2+^ responses to intracellular ROS generation in individual cells.

**Figure 2.**
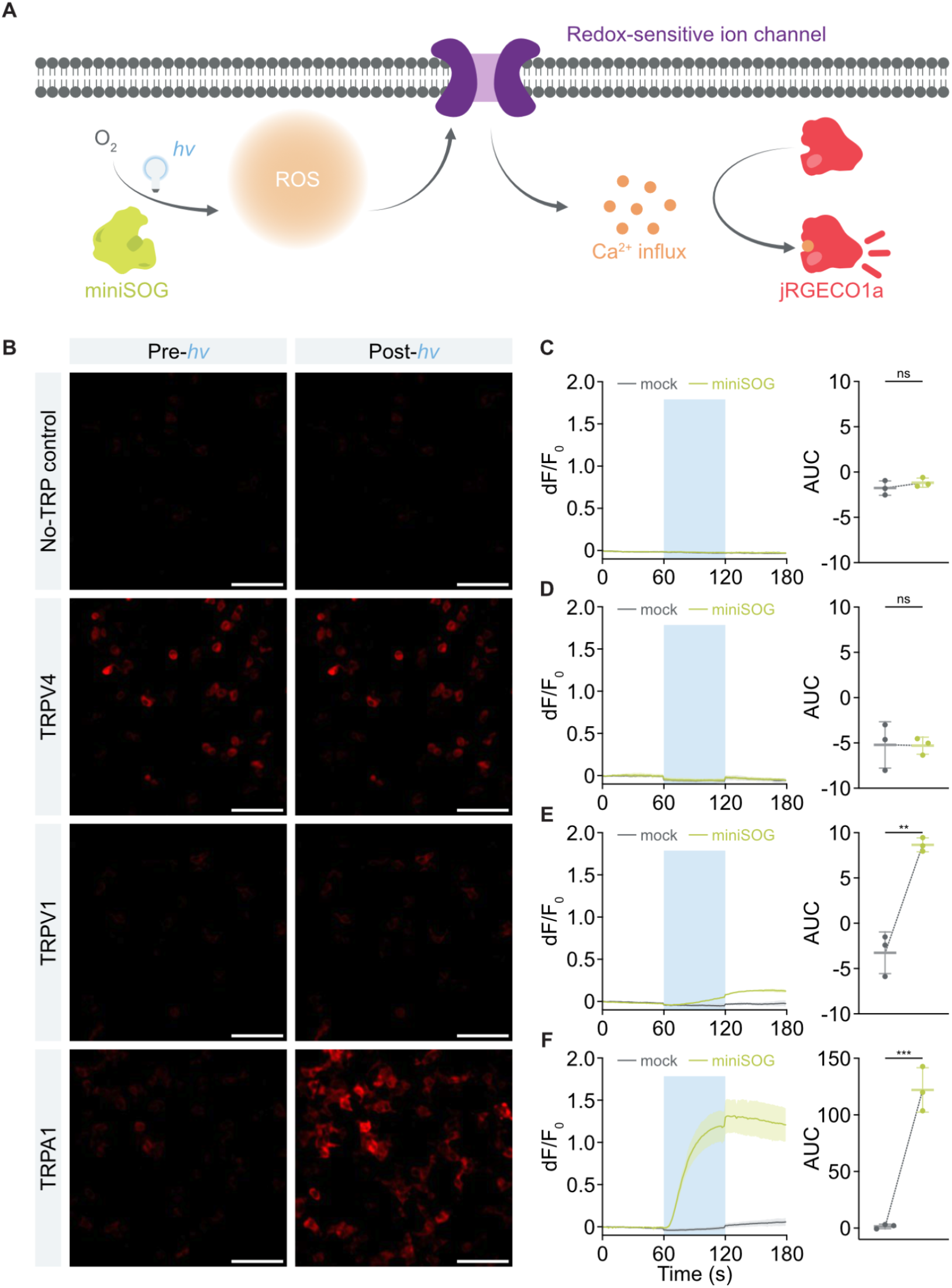
miniSOG-derived ROS activate redox-sensitive ion channels TRPV1 and TRPA1 in living cells. **(A)** Schematic of the experimental design. HEK293T cells were transiently transfected with miniSOG (or the corresponding empty vector for mock conditions), jRGECO1a, and either a target TRP channel (TRPV1, TRPV4, or TRPA1) or the corresponding empty vector for the no-TRP control condition. Blue-light illumination of miniSOG (470 nm, 60-120 s, 33 mW cm^−2^) generated ROS, while jRGECO1a fluorescence was monitored in real time to report intracellular Ca^2+^ dynamics. **(B)** Representative fluorescence images of jRGECO1a before and after blue-light stimulation for cells expressing the indicated channel. Scale bar, 100 μm. **(C-F)** Time-course traces (left) and corresponding AUC quantification (right) of jRGECO1a fluorescence for no-TRP control cells **(C)**, TRPV4-expressing cells **(D)**, TRPV1-expressing cells **(E)**, and TRPA1-expressing cells **(F)**. Blue shading indicates illumination period (60-120 s). Pronounced increases in ΔF/F_0_ were observed for TRPV1 and TRPA1 but not for TRPV4 or no-TRP control groups. Data are mean ± s.d. (n = 3 independent imaging dishes). Statistical significance was determined by two-tailed *t* tests; ns, not significant; **, *p* < 0.01; ***, *p* < 0.001.

Illumination elicited clear jRGECO1a fluorescence increases in TRPV1- and TRPA1-expressing cells, but not in TRPV4 or no-TRP control groups (**Figure 2B-F**). Importantly, TRPA1-expressing mock cells lacking miniSOG did not exhibit detectable Ca^2+^ responses upon blue-light illumination, indicating that the observed activation depends on miniSOG-mediated ROS generation rather than light alone. TRPA1 exhibited the largest overall response, whereas TRPV1 showed a smaller but reproducible activation relative to controls. Because responses were quantified as ΔF/F_0_ values normalized to each cell’s baseline fluorescence, differences in basal jRGECO1a intensity across conditions did not affect the measured activation amplitudes. Quantification of fluorescence AUCs confirmed significant activation of TRPV1 and TRPA1 in miniSOG-expressing cells compared with control groups (**Figure 2E,F**), whereas neither TRPV4 nor no-TRP control cells displayed measurable responses (**Figure 2C,D**). Further comparison between TRPV1 and TRPA1 revealed that TRPA1 reached both a higher peak amplitude and a shorter onset time (**Supplementary Figure 2**).

Together, these results demonstrate that genetically encoded ROS generation can functionally activate selected redox-sensitive ion channels in living cells and that this coupling is strongly target-dependent. The differential responsiveness of TRPV1, TRPV4, and TRPA1 indicates that identical ROS-generation conditions do not uniformly activate all membrane proteins, but instead reveal target-dependent differences in oxidative coupling. Notably, TRPA1 responses remained elevated after illumination ceased, indicating that oxidative processes initiated during the illumination period can sustain channel activation beyond the light window and that functional responses may persist transiently after the primary ROS input has ended. These differences prompted us to examine whether distinct oxidative pathways contribute differentially to modulation of soluble and membrane targets.

### Distinct ROS-dependent oxidative pathways underlie soluble and membrane target modulation

The selective responsiveness of TRPV1 and TRPA1, but not TRPV4, under identical illumination conditions prompted us to examine whether distinct oxidative pathways contribute differentially to target modulation. Although miniSOG was initially characterized as a singlet oxygen generator, subsequent work has shown that additional photochemical pathways can contribute to substrate oxidation^33^, raising the possibility that miniSOG-derived ROS engage multiple reactive pathways in cells. These observations suggest that miniSOG-derived oxidative effects may arise from multiple reactive pathways that couple differently to distinct molecular targets.

To probe the contribution of singlet oxygen-sensitive processes, we applied sodium azide (NaN_3_), a commonly used singlet oxygen quencher^34^. NaN_3_ significantly attenuated light-induced HyPerRed oxidation (**Figure 3A**), suggesting contribution from singlet oxygen-sensitive oxidation. Because HyPerRed derives from the OxyR redox-sensing module, which is responsive to small neutral oxidants in a solvent-accessible environment^23^, its attenuation by NaN_3_ is consistent with contribution from singlet oxygen-sensitive oxidation. TRPV1 activation was partially reduced by NaN_3_ (**Figure 3B**), whereas TRPA1 activation was largely unaffected (**Figure 3C**), indicating that distinct ROS-dependent oxidative pathways contribute to activation of the tested targets.

**Figure 3.**
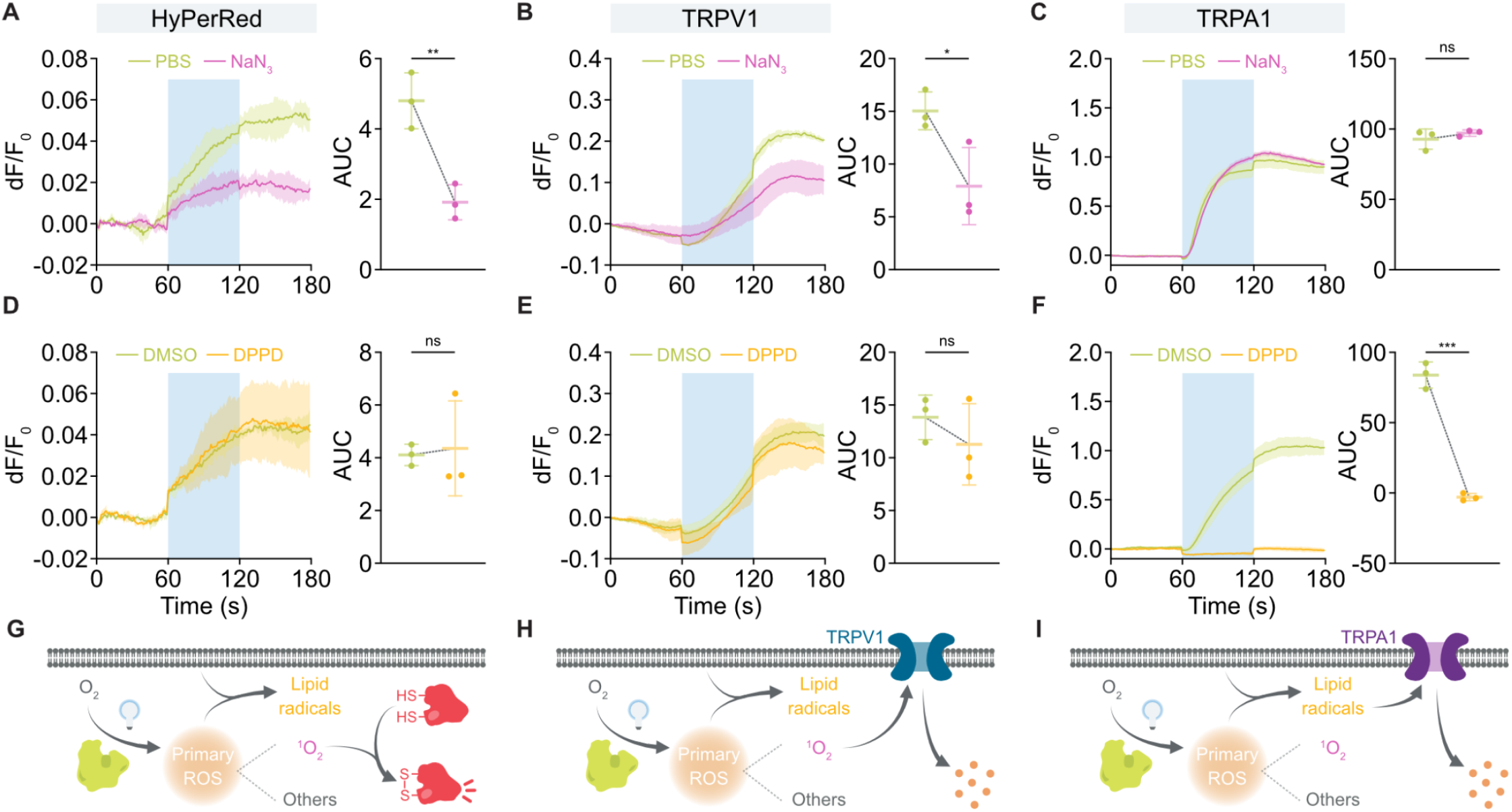
Distinct ROS-dependent oxidative pathways differentially contribute to soluble and membrane target modulation. **(A-C)** Time-course traces (left) and corresponding AUC quantification (right) of HyPerRed **(A)**, TRPV1 **(B)**, and TRPA1 **(C)** responses in the presence or absence of the singlet oxygen scavenger sodium azide (NaN_3_, 50 mM). Blue shading indicates illumination period (33 mW cm^−2^, 60 s). NaN_3_ attenuated HyPerRed and TRPV1 responses but not TRPA1 activation. **(D-F)** Time-course traces (left) and corresponding AUC quantification (right) of HyPerRed **(D)**, TRPV1 **(E)**, and TRPA1 **(F)** responses in the presence or absence of the lipid radical scavenger DPPD (25 μM). DPPD selectively suppressed TRPA1 activation, with no significant effect on HyPerRed or TRPV1 responses. Data are mean ± s.d. (n = 3 independent imaging dishes). Statistical significance was determined by two-tailed *t* tests; ns, not significant; *, *p* < 0.05; **, *p* < 0.01; ***, *p* < 0.001. **(G-I)** Conceptual models summarizing differential engagement of ROS in oxidative modulation of HyPerRed **(G)**, TRPV1 **(H)**, and TRPA1 **(I)**. Blue-light excitation of miniSOG generates a mixture of primary ROS, including singlet oxygen (^1^O_2_) and other oxidants. In membrane environments, primary ROS can additionally initiate lipid peroxidation, leading to lipid radical formation. Functional modulation of individual targets reflects preferential coupling to distinct ROS-dependent chemistries.

We next examined the involvement of membrane-associated oxidative processes using N,N’-diphenyl-p-phenylenediamine (DPPD), a lipid-phase radical scavenger that inhibits lipid peroxidation. DPPD had minimal effects on HyPerRed fluorescence or TRPV1 activation (**Figure 3D, E**), but strongly suppressed TRPA1 activation (**Figure 3F**). This complementary selectivity compared with NaN_3_ suggests that membrane-associated lipid radical processes play a particularly important role in TRPA1 modulation. This differential sensitivity suggests that distinct ROS-dependent oxidative pathways preferentially couple to different protein targets. Together, these results indicate that miniSOG-generated ROS engage multiple oxidative pathways in cells and that individual targets differ in how strongly they couple to those pathways. Soluble redox reporters and membrane ion channels exhibit distinct sensitivities to pharmacological scavengers, indicating that oxidative signaling outcomes depend on both ROS chemistry and the local molecular environment (**Figure 3G-I**).

### Genetically encoded oxidative modulation of TRPA1 is tunable by illumination and expression parameters

Having identified TRPA1 as the most responsive channel to miniSOG-derived ROS, we next asked whether its activation could be quantitatively tuned by optical and genetic parameters. Because the extent of oxidative modulation depends on both the amount of ROS produced and the relative abundance between the photosensitizer and the channel, we systematically varied light intensity, light duration, and the mass ratio between the genetic constructs of TRPA1 and miniSOG, thereby establishing key optical and genetic parameters that tune activation efficiency (**Figure 4**).

**Figure 4.**
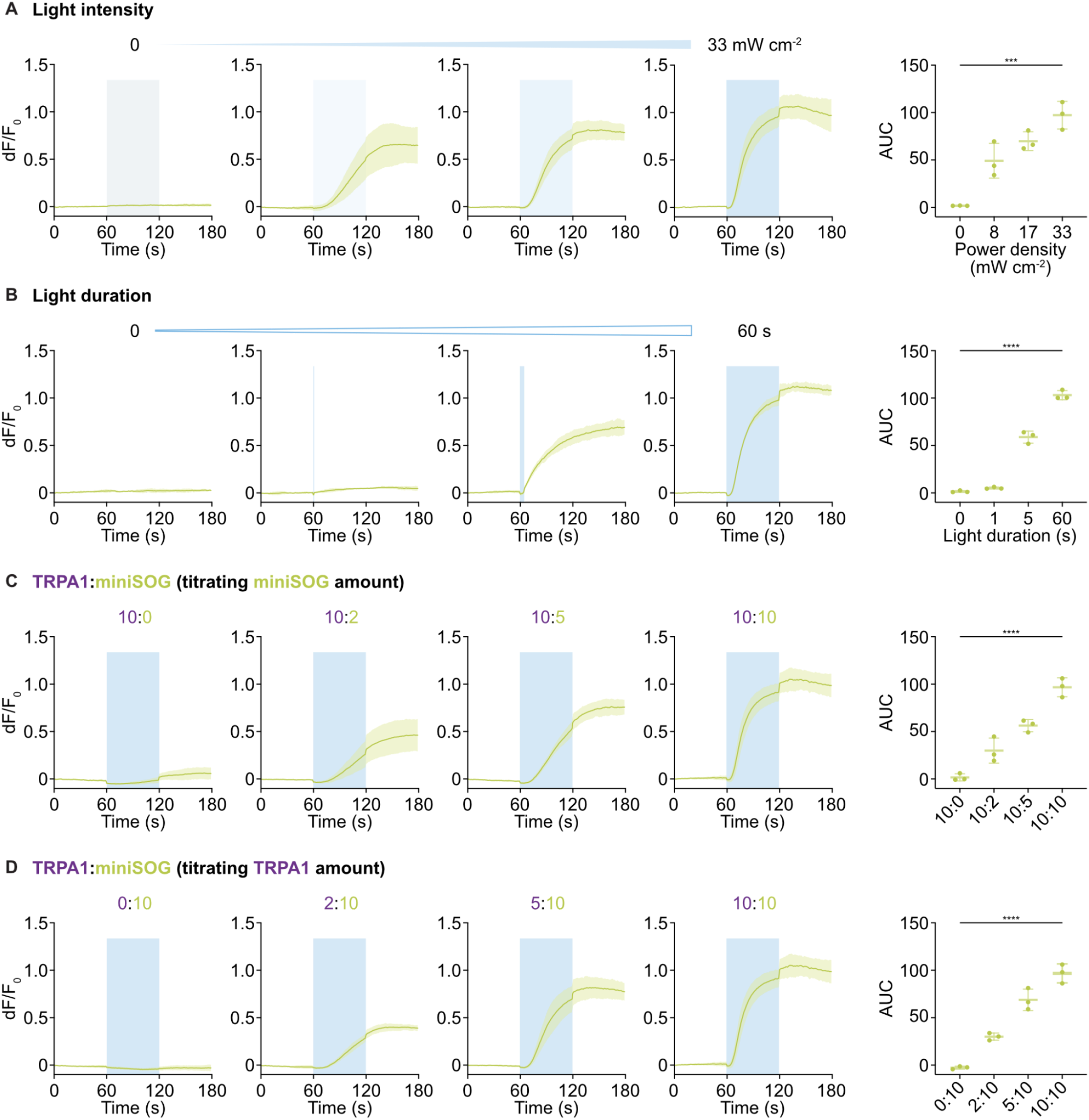
Optical and genetic parameters tune TRPA1 activation by miniSOG-derived ROS. **(A)** Effect of light intensity on TRPA1 activation. Blue-light illumination (60-120 s) at increasing power densities produced progressively larger Ca^2+^ responses in HEK293T cells co-expressing miniSOG and TRPA1, whereas no change occurred at 0 mW cm^−2^. Quantification of the AUCs showed intensity-dependent activation. **(B)** Effect of light duration at a fixed intensity of 33 mW cm^−2^. Short 1 s stimulation elicited minimal responses, whereas longer exposures (5 s and 60 s) produced progressively larger responses. Corresponding AUC quantification showed duration-dependent increases. **(C-D)** Effect of relative expression of TRPA1 and miniSOG on activation. Increasing the plasmid mass of either miniSOG **(C)** or TRPA1 **(D)** enhanced Ca^2+^ responses, consistent with dependence on both ROS source and target abundance. Data are mean ± s.d. (n = 3 independent imaging dishes). Statistical significance was determined by one-way ANOVA with multiple comparisons; ***, *p* < 0.001; ****, *p* < 0.0001.

Increasing light intensity led to progressively larger Ca^2+^ transients in TRPA1-expressing cells co-expressing miniSOG, whereas no detectable change occurred at 0 mW cm^−2^ (**Figure 4A**). Quantification of the integrated fluorescence response (AUC) increased monotonically with light intensity, consistent with greater oxidative input. However, the peak ΔF/F_0_ did not differ significantly across intensities, while the activation onset time shortened at higher intensities (**Supplementary Figure 3A**). These results indicate that stronger illumination primarily accelerates TRPA1 activation kinetics and modestly enhances the overall integrated response, rather than increasing the peak amplitude of Ca^2+^ influx.

Increasing light duration at a fixed intensity of 33 mW cm^−2^ also led to progressively stronger TRPA1 activation (**Figure 4B**). Brief 1 s stimulation triggered small Ca^2+^ responses, whereas 5 s stimulation markedly enhanced activation, and 60 s exposure yielded the largest signal. Quantification showed that both the AUC and the peak ΔF/F_0_ increased with longer light duration, while the onset time decreased significantly (**Supplementary Figure 3B**). No detectable response was observed in the 0 s control, confirming that activation required photoexcitation of miniSOG. These results demonstrate that TRPA1 activation scales with illumination duration primarily through an increase in response magnitude and accelerated activation kinetics.

Increasing the plasmid mass of either miniSOG or TRPA1 enhanced the Ca^2+^ response (**Figure 4C, D**). When TRPA1 plasmid mass was held constant, raising miniSOG plasmid mass produced progressively stronger and faster Ca^2+^ responses, as shown by increased AUC, higher peak ΔF/F_0_, and shorter onset time (**Supplementary Figure 3C**,**D**). Conversely, when miniSOG was fixed, increasing TRPA1 plasmid mass likewise strengthened activation and accelerated response kinetics. These results demonstrate that both the oxidant source and its target contribute to the overall efficiency of genetically encoded oxidative modulation, and that oxidative control of TRPA1 scales with the absolute expression levels of both components.

To directly assess whether a representative HEK293T operating condition used for oxidative modulation causes overt compromise of cellular integrity, we next evaluated cell viability under the highest miniSOG DNA input and illumination condition used in the HEK293T parameterization experiments. Specifically, HEK293T cells expressing miniSOG alone at the highest miniSOG DNA input tested above, without co-expression of additional effector or reporter proteins, were exposed to blue light at 33 mW cm^−2^ for 60 s. Flow cytometric analysis using SYTOX Blue showed no detectable reduction in viability at either 4 h or 24 h post-stimulation, and no significant difference in cell number was observed between illuminated and non-illuminated cells at 24 h (**Supplementary Figures 4, 5, and 6**). Together, these data indicate that miniSOG expression and optical stimulation under this representative HEK293T upper-bound condition do not cause detectable overt loss of cellular viability or cell number in HEK293T cells. These assays do not exclude all possible sublethal biochemical effects of ROS generation, but they support the conclusion that miniSOG expression and optical stimulation do not produce detectable overt cytotoxicity or short-term cell loss under the representative HEK293T operating condition tested here. Having defined a tunable operating regime, we next asked whether oxidative modulation could be further organized through spatial positioning of the ROS source.

### Membrane localization of miniSOG enhances oxidative modulation of TRPA1

To determine whether subcellular positioning of the ROS source influences the efficiency of oxidative modulation, we examined how membrane localization of miniSOG affects TRPA1 activation. In previous experiments, cytosolic miniSOG generated ROS throughout the cell, which may limit oxidative reactions within the plasma membrane lipid environment where TRPA1 resides. We therefore compared cytosolic and membrane-targeted (LCK-tagged) miniSOG constructs co-expressed with TRPA1 in HEK293T cells under identical illumination conditions (**Figure 5A**).

**Figure 5.**
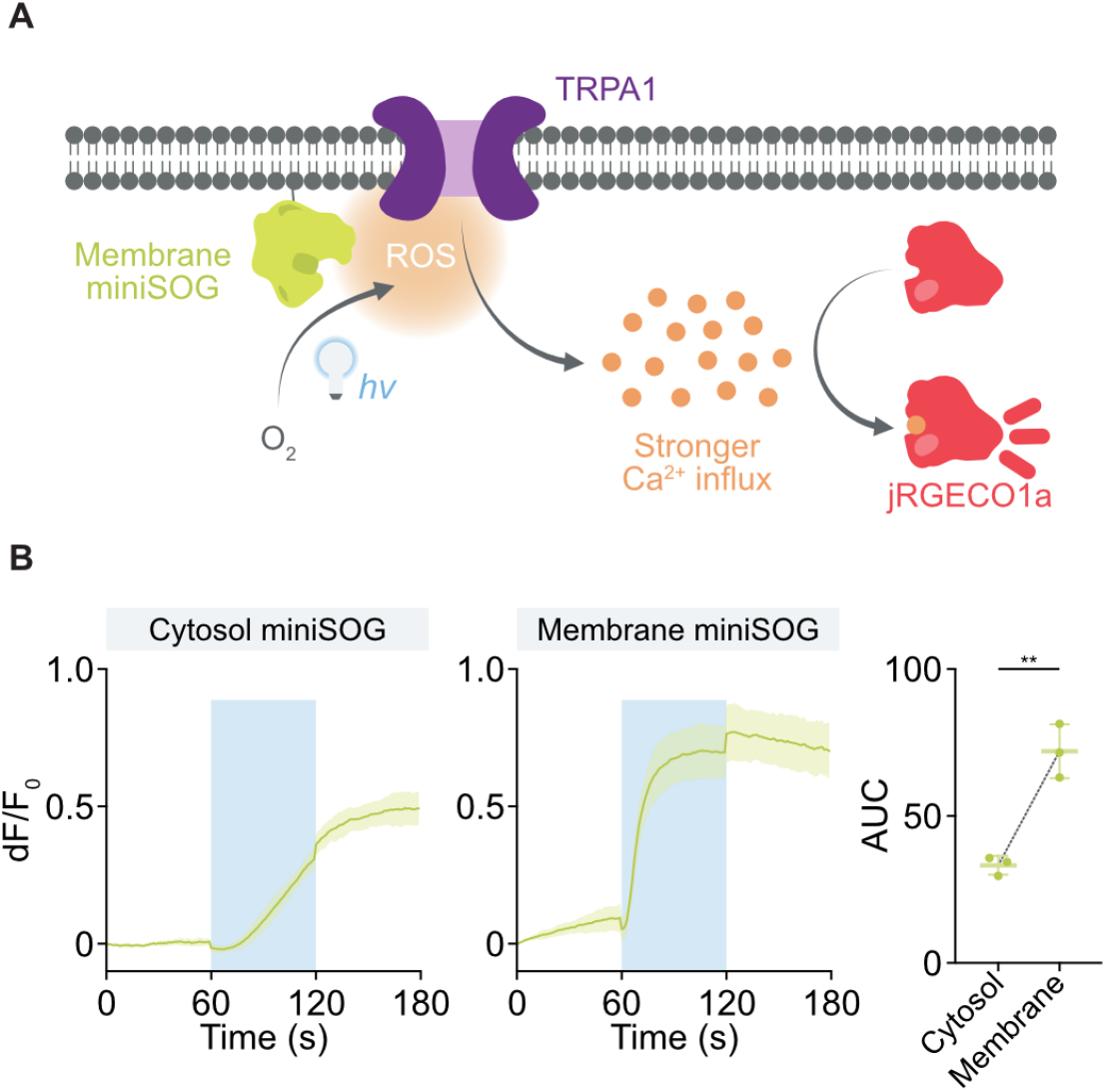
Membrane localization of miniSOG enhances oxidative modulation of TRPA1. **(A)** Schematic illustrating enhanced TRPA1 activation when miniSOG is anchored to the plasma membrane. Blue-light illumination of membrane-targeted miniSOG generates ROS in closer proximity to the channel, promoting stronger Ca^2+^ influx detected by jRGECO1a. **(B)** Time-course traces (left) and corresponding AUC quantification (right) of TRPA1 activation in HEK293T cells co-expressing cytosolic or membrane-targeted miniSOG under identical illumination conditions (33 mW cm^−2^, 60-120 s). Membrane-anchored miniSOG produced a significantly larger integrated fluorescence response. Data are mean ± s.d. (n = 3 independent imaging dishes). Statistical significance was determined by two-tailed *t* tests; **, *p* < 0.01.

Illumination from 60-120 s induced a pronounced fluorescence increase in TRPA1-expressing cells with membrane-targeted miniSOG, whereas the cytosolic form elicited a weaker response (**Figure 5B**). Quantification showed that membrane localization increased the peak ΔF/F_0_ and accelerated activation kinetics, reflected by a shorter onset time (**Supplementary Figure 7**). The integrated AUC also increased, consistent with more efficient oxidative modulation when the ROS generator is positioned near the membrane lipid environment.

To confirm that these responses required TRPA1, we expressed soluble or membrane-targeted miniSOG alone under identical illumination conditions. Neither construct produced detectable Ca^2+^ signals in the absence of TRPA1 (**Supplementary Figure 8**), indicating that the observed activation arose from oxidation-dependent gating of the channel rather than direct photostimulation or secondary signaling. Together, these findings show that subcellular positioning of the oxidant source influences oxidative modulation, motivating us to explore strategies that further bias ROS generation toward the molecular neighborhood of a target protein.

### Fusion-based targeting further biases oxidative modulation toward the molecular neighborhood of target proteins

Going a step beyond subcellular targeting, we asked whether fusion-based targeting, an approach commonly used in CALI^6,8,9^, could be incorporated into our framework to provide an additional level of spatial constraints. To test this possibility, we fused miniSOG to TRPV1, which preliminary experiments showed to be amenable to N-terminal miniSOG fusion.

Specifically, we generated a miniSOG-GGGS-TRPV1 fusion construct (**Figure 6A**). Under blue-light illumination, HEK293T cells expressing the miniSOG-GGGS-TRPV1 fusion exhibited a light-evoked increase in intracellular Ca^2+^, whereas cells expressing TRPV1 showed no detectable response under identical conditions (**Figure 6B**). Quantification of the integrated fluorescence response confirmed activation in the fusion condition. These results show that fusion-based targeting can be implemented within genetically encoded oxidative modulation and provides a practical route to bias ROS generation toward the molecular neighborhood of a protein of interest. Together with the membrane-localization experiments (**Figure 5**), these results support a multi-scale view in which oxidative modulation can be spatially biased at both subcellular and protein-proximal scales, without requiring absolute molecular confinement of the oxidative signal.

**Figure 6.**
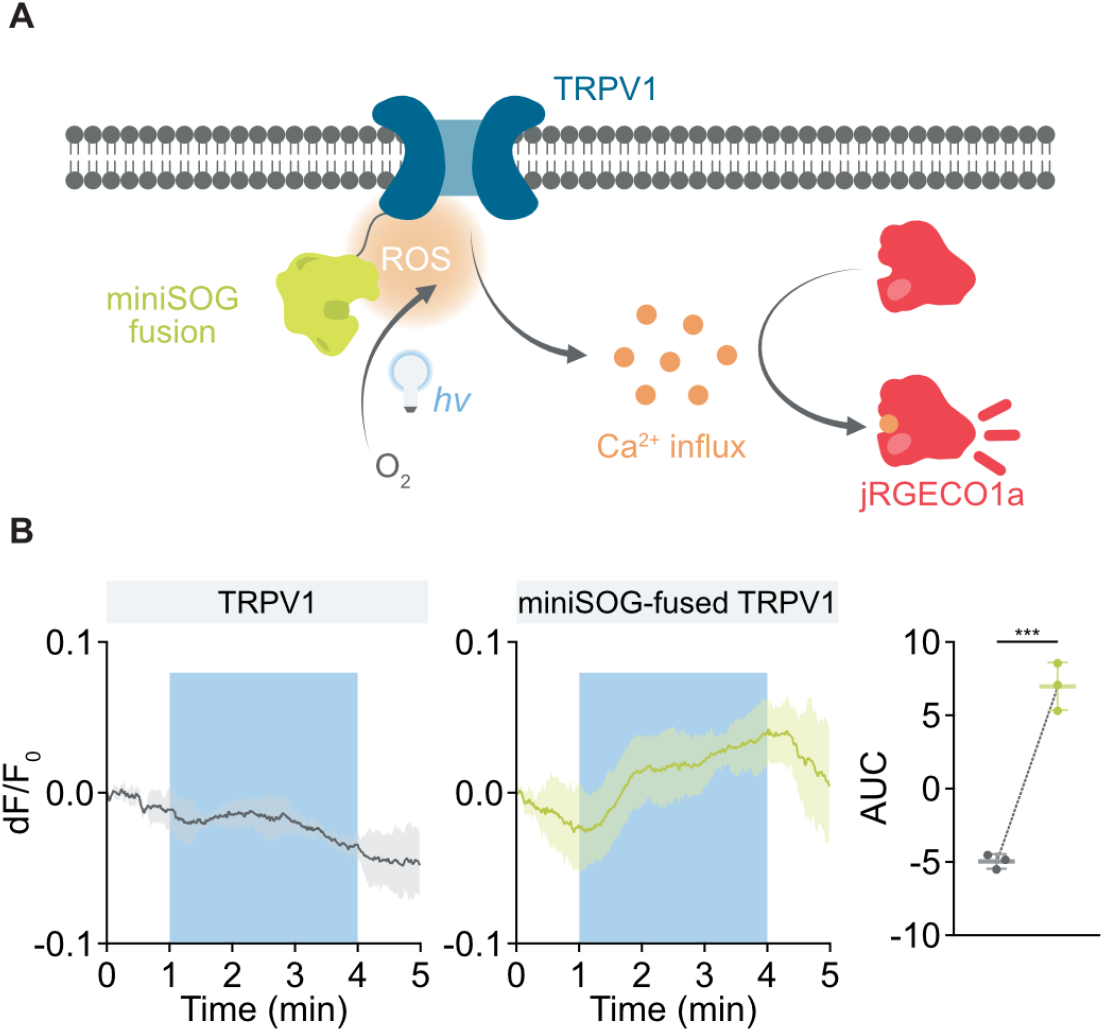
Fusion-based targeting enables protein-proximal oxidative modulation of TRPV1. **(A)** Schematic illustrating protein-proximal ROS generation through a miniSOG-GGGS-TRPV1 fusion construct. Blue-light illumination of the fused miniSOG generates ROS in close proximity to TRPV1, promoting TRPV1 activation and Ca^2+^ influx detected by jRGECO1a. **(B)** Time-course traces (left) and corresponding AUC quantification (right) of TRPV1 activation in HEK293T cells expressing TRPV1 or the miniSOG-GGGS-TRPV1 fusion under identical illumination (66 mW cm^−2^, 1-4 min). The fusion construct produced a robust integrated fluorescence response, whereas TRPV1 alone showed no detectable activation under the same illumination conditions. Data are mean ± s.d. (n = 3 independent imaging dishes). Statistical significance was determined by two-tailed *t* tests; ***, *p* < 0.001.

### Genetically encoded ROS activate endogenous TRPA1 channels in human fibroblasts

Having shown that genetically encoded ROS can modulate heterologously expressed TRP channels, we next asked whether this strategy could also regulate an endogenously expressed target in its native cellular context. We therefore performed experiments in IMR-90 human fibroblasts, a cell type reported to express endogenous TRPA1 channels^35–37^. Cells were co-transfected with miniSOG and jRGECO1a and subjected to blue-light illumination at 66 mW cm^−2^ for 3 min (**Figure 7A**).

**Figure 7.**
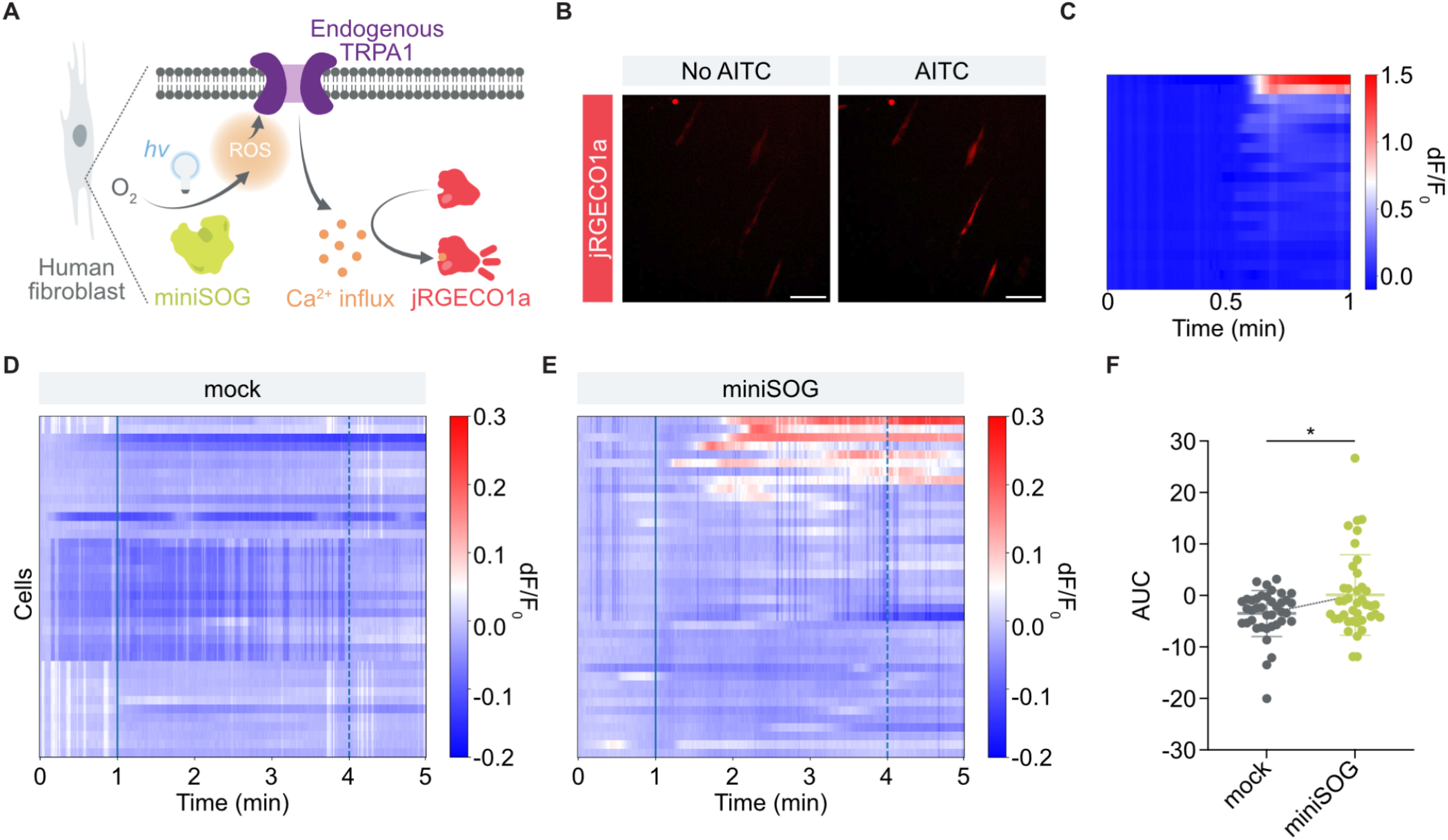
Genetically encoded ROS generation activates endogenous TRPA1 channels in human fibroblasts. **(A)** Schematic illustrating activation of endogenous TRPA1 in IMR-90 human fibroblasts by miniSOG-generated ROS. Blue-light illumination of miniSOG produces ROS that stimulate TRPA1 channels, leading to Ca^2+^ influx detected by the fluorescent indicator jRGECO1a. **(B)** Representative fluorescence images of jRGECO1a responses in IMR-90 cells in the absence or presence of the TRPA1 agonist AITC (200 μM). Scale bar, 100 μm. **(C)** Heatmap showing single-cell jRGECO1a fluorescence response in IMR-90 cells upon stimulation with the TRPA1 agonist AITC (200 μM). Only a subset of cells exhibited strong Ca^2+^ responses, consistent with heterogeneous TRPA1 expression in this cell population. These data confirm the presence of functional TRPA1 channels in IMR-90 cells. **(D**,**E)** Single-cell Ca^2+^ response heatmaps for mock-transfected cells **(D)** and miniSOG-expressing cells **(E)** during blue-light illumination (66 mW cm^−2^, 1-4 min). **(F)** Quantification of Ca^2+^ responses shown as integrated fluorescence (AUC) for mock and miniSOG conditions. Data are mean ± s.d. (mock: n = 39 single cells; miniSOG: n = 40 single cells). Statistical significance was determined by two-tailed *t* tests; *, *p* < 0.05.

To first verify the presence of functional TRPA1 channels in IMR-90 cells, we applied the TRPA1 agonist allyl isothiocyanate (AITC, 200 μM). A subset of cells exhibited robust increases in jRGECO1a fluorescence following AITC stimulation (**Figure 7B-C**), indicating that functional TRPA1 channels are present but heterogeneously expressed in this cell population.

Blue-light illumination of miniSOG elicited increases in intracellular Ca^2+^ levels in a subset of IMR-90 cells (**Figure 7E**), consistent with the heterogeneous TRPA1 expression observed in the AITC validation experiments. In contrast, cells transfected with an empty vector control (mock) showed no detectable response under identical conditions (**Figure 7D**). Quantification of integrated fluorescence signals confirmed significantly larger Ca^2+^ responses in miniSOG-expressing cells (**Figure 7F**).

To determine whether these responses were mediated by endogenous TRPA1 channels, we performed pharmacological inhibition experiments using the selective TRPA1 antagonist HC-030031. Inhibitor treatment markedly suppressed the light-evoked Ca^2+^ responses in miniSOG-expressing IMR-90 cells (**Supplementary Figure 10**), confirming that the observed activation depends on TRPA1 activity. Together, these results demonstrate that genetically encoded ROS generation can modulate an endogenously expressed ion channel in its native cellular environment, thereby extending oxidative modulation beyond heterologous expression systems. In this framework, the IMR-90 experiments establish native endogenous-target modulation, while the controlled heterologous-expression experiments above define tunability and subcellular localization as design parameters for oxidative modulation.

## Discussion

Genetically encoded oxidative chemistry provides a tunable framework for modulating protein function in living cells. By co-expressing the photosensitizer miniSOG with distinct protein targets, we demonstrate that intracellular ROS generation can alter protein activity through oxidative chemistry across multiple protein classes. In the soluble redox reporter, miniSOG-derived ROS shifted the redox equilibrium and increased HyPerRed fluorescence, whereas in membrane ion channels such as TRPA1 and TRPV1, ROS generation triggered redox-dependent Ca^2+^ influx. Together, these findings support the view that genetically encoded ROS production can serve as a versatile means to functionally modulate both soluble and membrane proteins. Importantly, we further demonstrate that this strategy can regulate an endogenously expressed ion channel in human fibroblasts, indicating that genetically encoded oxidative chemistry can operate within native cellular signaling contexts. In contrast to prior applications of genetically encoded photosensitizers that primarily exploit ROS for CALI, labeling, or phototoxic ablation^22^, our results highlight their use as programmable chemical inputs for regulating protein function. Because ROS-triggered oxidation can initiate secondary chemical and signaling cascades, functional responses can outlast the illumination window even after ROS generation ceases. Temporal control in this system is therefore exerted at the level of initiating oxidative input rather than necessarily at the level of immediate termination of all downstream consequences.

Consistent with this view, our pharmacological scavenger experiments suggest that different targets preferentially couple to distinct oxidative pathways. Quenching singlet oxygen with sodium azide attenuated HyPerRed oxidation and partially reduced TRPV1 activation, whereas TRPA1 activation was largely insensitive to singlet oxygen scavenging but strongly suppressed by the lipid radical scavenger DPPD. These observations indicate that miniSOG-generated ROS can engage multiple downstream oxidative chemistries in cells, including singlet-oxygen-mediated reactions and membrane-associated lipid radical processes. While these experiments do not resolve the precise molecular oxidation events at each target, the differential sensitivity to pathway-selective scavengers indicates that functional modulation arises from target-dependent coupling to distinct ROS-driven oxidative processes rather than from a single nonspecific oxidative cascade. The functional outcome therefore reflects how individual protein targets couple to these distinct oxidative pathways within their local molecular environment. This target-dependent coupling to distinct oxidative pathways suggests a chemical basis for selectivity-by-context, a key consideration for designing oxidative regulation.

Systematic variation of optical and genetic parameters reveals that oxidative modulation is quantitatively tunable. TRPA1 activation scales with illumination intensity, duration, and the relative expression ratio of miniSOG and TRPA1, defining a tunable parameter space for ROS-based regulation. Consistent with this defined operating regime, miniSOG expression and optical stimulation under a representative HEK293T upper-bound condition did not cause detectable overt loss of cell viability or cell number, supporting the feasibility of implementing oxidative modulation under this tested HEK293T operating condition. These data should not be interpreted as showing that miniSOG activation is globally inert or free of all sublethal oxidative biochemical effects. This bounded interpretation is consistent with the use of ROS-generating and photosensitizer-based approaches in living systems^9,38,39^, where claims are supported by defined operating conditions and mechanism-relevant functional controls rather than by exhaustive exclusion of all possible downstream biochemical effects. Rather, together with the target-dependent TRP responses, pathway-selective scavenger effects, no-TRP controls, and pharmacological inhibition in IMR-90 cells, these data support the conclusion that the functional outputs observed here are not explained by overt cytotoxicity or generalized cellular failure. These results indicate that oxidative modulation can be adjusted in a graded manner through optical regulation of ROS generation. Such quantitative control positions engineered ROS generation as a programmable regulatory layer that can be tuned independently through optical and genetic inputs.

Our results further highlight the importance of spatial organization in implementing oxidative modulation. Membrane anchoring of miniSOG first establishes that subcellular positioning of the ROS source influences modulation efficiency, whereas fusion-based targeting extends this principle by introducing an additional protein-proximal layer for biasing oxidative modulation. Importantly, because ROS are inherently reactive and diffusible, these approaches are best viewed as biasing oxidative coupling toward defined targets through localization, rather than requiring absolute molecular confinement. This interpretation is consistent with established CALI and proximity-labeling applications of genetically encoded photosensitizers, in which photosensitizer-derived ROS have been used to preferentially perturb or label nearby biomolecules in a localization-dependent manner^6,9,18,22^. Together, the subcellular and protein-proximal implementations support a multi-scale view in which oxidative modulation can be organized at progressively finer spatial scales. At the same time, the present study does not directly resolve the molecular spatial range or decay kinetics of the oxidative species and their secondary downstream chemistries, and therefore the system is best viewed as spatially biased rather than strictly confined in space and time.

More broadly, this work reframes oxidative chemistry, traditionally associated with cellular stress, as a programmable regulatory mechanism when deployed in a genetically encoded and quantitatively controlled manner. Conceptually, this work extends genetically encoded photosensitizers beyond their established roles in phototoxicity and CALI to a versatile strategy for engineering protein regulation using localized oxidative chemistry. By combining tunable ROS generation with modular localization strategies, genetically encoded oxidative chemistry can be integrated into synthetic biology toolkits as a flexible platform for redox-dependent regulation. The ability to modulate both engineered and endogenous targets highlights its potential as a general strategy for manipulating protein function in living cells. Future efforts may further characterize the spatial range, temporal decay, and molecular selectivity of engineered ROS signals, explore additional ROS-sensitive effectors, and incorporate complementary biochemical or proteomic analyses to assess target specificity. Rather than aiming for absolute molecular precision, these directions position genetically encoded oxidative chemistry as a programmable and spatially structured approach for modulating protein function in complex cellular environments.

## Methods

### Plasmid construction and molecular biology

All plasmids were assembled using Gibson Assembly (New England Biolabs, NEB) from PCR fragments generated with Q5 High Fidelity DNA polymerase (NEB). Assemblies were transformed into NEB Stable *E. coli* by heat shock for plasmid propagation, and all constructs were verified by commercial whole-plasmid sequencing (Plasmidsaurus, USA).

The plasmid containing the coding sequence of HyPerRed, pC1-HyPerRed, was a gift from Vsevolod Belousov (Addgene plasmid # 48249; http://n2t.net/addgene:48249; RRID:Addgene_48249). Plasmids encoding miniSOG (pCMVSp-miniSOG), jRGECO1a (pCMVSp-jRGECO1a), mouse TRPA1 (pCMV-mTRPA1), rat TRPV1 (pCMV-rTRPV1), and rat TRPV4 (pCMV-rTRPV4) were generated in-house using PCR amplification and Gibson Assembly. The miniSOG-GGGS-TRPV1 fusion construct (pCMVSp-miniSOG-GGGS-TRPV1) was generated by inserting the miniSOG coding sequence upstream of rat TRPV1 with a flexible N-terminal GGGS linker using Gibson Assembly. PCR primers were synthesized by Integrated DNA Technologies (IDT). For mock control experiments, the empty vector corresponding to the miniSOG expression plasmid was used to maintain equal total DNA mass during co-transfection. For no-TRP control experiments, the empty vector corresponding to the channel expression plasmid was used to maintain equal total DNA mass during co-transfection.

### HEK293T cell culture and transient transfection

HEK293T cells (American Type Culture Collection (ATCC), CRL-3216) were cultured in Dulbecco’s Modified Eagle Medium (DMEM, Corning, 10-013-CV) supplemented with 10% fetal bovine serum (FBS, Gibco) and 1× penicillin-streptomycin at 37°C and 5% CO_2_ in a humidified incubator. Cells were used at passages below 20. For imaging experiments, cells were seeded onto 14-mm glass-bottom dishes (Avantor, D35-14) pre-coated with bovine collagen to promote cell adhesion. Glass-bottom dishes were coated with 200 μL of bovine collagen solution (100 μg mL^−1^ in 10 mM HCl, prepared from a 10 mg mL^−1^ telocollagen stock, CELLINK) for 3 h at room temperature. Dishes were then rinsed with phosphate-buffered saline (PBS, Cytiva HyClone, SH30028.02) prior to cell seeding. Cells were detached using 0.25% trypsin-EDTA (Thermo Fisher Scientific, 25200056), centrifuged at 500 × g for 5 min, and resuspended in complete culture medium. Approximately 1.6 × 10^5^ cells were seeded per dish in 200 μL medium and allowed to attach for 3 h, after which an additional 300 μL medium was added. Cells were cultured overnight prior to transfection.

Cells were transfected at approximately 80% confluency. Transient transfection mixtures were prepared using polyethyleneimine (PEI-MAX, Polysciences) at a PEI:DNA mass ratio of 2.58:1. Plasmids were mixed at the indicated mass ratios prior to complex formation. The mixture was incubated for 12 minutes at room temperature and added dropwise to the cells.

For HyPerRed reporter experiments, cells were co-transfected with 160 ng pC1-HyPerRed and 160 ng pCMVSp-miniSOG per dish (320 ng total DNA). In mock control conditions, 160 ng HyPerRed plasmid was co-transfected with 160 ng of the parental corresponding empty vector used for the miniSOG construct to maintain equal DNA mass. For ion channel co-expression experiments, cells were co-transfected with 110 ng channel plasmid (or empty vector for no channel controls), 110 ng pCMVSp-miniSOG (or empty vector for mock controls), and 110 ng pCMVSp-jRGECO1a per dish (330 ng total DNA), maintaining equal DNA mass across conditions. For miniSOG-GGGS-TRPV1 fusion experiments, cells were co-transfected with 160 ng TRPV1 or miniSOG-GGGS-TRPV1 plasmid together with 160 ng jRGECO1a plasmid per dish (320 ng total DNA). Fresh culture medium was added 24 h after transfection to reach a final volume of 3.5 mL prior to imaging.

### IMR-90 cell culture and transient transfection

Human lung fibroblast IMR-90 cells (ATCC, CCL-186) were cultured in DMEM (Thermo Fisher Scientific, 11965092) supplemented with 10% FBS and 1× MEM non-essential amino acids (Thermo Fisher Scientific, 11140050) at 37°C and 5% CO_2_ in a humidified incubator. Cells between passages 2 and 5 were used for experiments. For imaging experiments, cells were seeded onto 14-mm glass-bottom dishes pre-coated with bovine collagen to promote fibroblast adhesion. Glass-bottom dishes were coated with 200 μL of bovine collagen solution (100 μg mL^−1^ in 10 mM HCl, prepared from a 10 mg mL^−1^ telocollagen stock, CELLINK) for 3 h at room temperature. Dishes were then rinsed with PBS prior to cell seeding. Cells were detached using 0.25% trypsin-EDTA, centrifuged at 125 × g for 10 min, and resuspended in complete culture medium. Approximately 4 × 10^4^ cells were seeded per dish in 200 μL medium and allowed to attach for 3 h, after which an additional 300 μL medium was added. Cells were cultured overnight prior to transfection.

Cells were transfected at approximately 80% confluency. Transient transfection was performed using Lipofectamine LTX and PLUS reagents (Thermo Fisher Scientific, A12621). For each dish, 500 ng total plasmid DNA was diluted in 100 μL Opti-MEM reduced-serum medium (Thermo Fisher Scientific, 31985062). DNA was mixed with 0.5 μL PLUS reagent and incubated for 10 min at room temperature. Lipofectamine LTX (1.25 μL) was then added and the mixture incubated for an additional 25 min before being added dropwise to the cells.

For IMR-90 calcium imaging experiments, cells were co-transfected with 250 ng pCMVSp-miniSOG (or the corresponding empty vector for mock control conditions) and 250 ng pCMVSp-jRGECO1a per dish (500 ng total DNA), maintaining equal DNA mass across conditions. Lipofectamine-based transfection was used for IMR-90 cells because these primary fibroblasts exhibited low transfection efficiency with PEI-based methods.

### Chemical treatment

The singlet oxygen scavenger sodium azide (NaN_3_, Sigma-Aldrich, S2002) and the lipid radical scavenger N,N’-diphenyl-p-phenylenediamine (DPPD, Sigma-Aldrich, 292265) were used at the concentrations indicated in the figure legends. NaN_3_ was prepared as a concentrated stock solution in PBS and diluted directly into the culture medium to the indicated final concentrations prior to imaging. DPPD was prepared as a stock solution in dimethyl sulfoxide (DMSO, Thermo Fisher Scientific, D12345) and diluted into the imaging medium to the indicated final concentrations prior to experiments. For vehicle control conditions, an equivalent volume of solvent was added.

For TRPA1 inhibition experiments in IMR-90 cells, the selective TRPA1 antagonist HC-030031 (MedChemExpress, HY-10012) was prepared as a stock solution in DMSO and diluted into the culture medium to the indicated final concentration prior to imaging. Cells were incubated with HC-030031 for 30 min before optical stimulation to allow inhibition of TRPA1-mediated calcium influx.

### Fluorescent imaging and optical stimulation

A 575 nm LED light source (Lumencor) was used to excite HyPerRed and jRGECO1a, and emitted signals were collected through a 10× objective (NA 0.3, Leica) and recorded using an sCMOS camera (Zyla 5.5, Andor) at 2 Hz. A 470 nm LED light source (Lumencor) was used to stimulate miniSOG at 2 Hz with an exposure duration of 400 ms per frame. The blue-light power densities for stimulation are indicated in the corresponding figure legends. Blue-light pulses were synchronized with image acquisition. Because the multipass dichroic filter set used in the imaging path (Chroma 89402bs) transmits miniSOG fluorescence, frames coinciding with blue-light illumination contained detectable miniSOG signals and were excluded from analysis. Consequently, every second frame corresponding to blue-light illumination was removed prior to analysis, and the remaining frames acquired between stimulation pulses were used for downstream quantification. Cells were imaged in complete culture medium while the culture dish was placed in a 37°C water bath to maintain physiological temperature during optical stimulation. Recorded images were processed to extract HyPerRed or calcium signals from individual cells. For HEK293T experiments, individual cell bodies were automatically segmented using CellPose^40^, typically identifying hundreds to thousands of cells per dish. For IMR-90 experiments, which typically contained relatively few jRGECO1a-positive cells per field of view, individual cells were manually defined as regions of interest (ROIs). ROIs were randomly selected from jRGECO1a-expressing cells within each field of view, without regard to their calcium response, and fluorescence intensity traces were extracted for each cell.

For most HEK293T experiments, blue-light stimulation was applied between 60 s and 120 s of the recording. For miniSOG-GGGS-TRPV1 fusion experiments, the stimulation period was extended from 60 s to 240 s. For IMR-90 calcium imaging experiments, blue-light stimulation was applied between 60 s and 240 s of the recording.

### Cell viability analysis

For cell viability experiments, HEK293T cells were seeded and transfected as described above. Each sample was transfected with 110 ng pCMVSp-miniSOG together with 220 ng of the corresponding parental empty vector, for a total DNA mass of 330 ng per dish. Blue-light stimulation was performed under the same optical conditions described above. After stimulation, cells were returned to the incubator and collected at either 4 h or 24 h post-stimulation for viability analysis. Culture medium was removed, and cells were dissociated by adding 200 μL 0.25% trypsin-EDTA per dish. After dissociation, trypsin was neutralized with 200 μL of complete culture medium. Cells were collected by centrifugation at 500 × g for 5 min, the supernatant was removed, and the cell pellet was resuspended in PBS. SYTOX Blue dead cell stain (Invitrogen, S34857) was then added, and cells were incubated for 5 min at room temperature before flow cytometry analysis. Flow cytometry data were analyzed using the gating strategy shown in Supplementary Figure 4. Cell viability was defined as the fraction of SYTOX Blue-negative cells, and cell number at 24 h was quantified as the number of SYTOX Blue-negative events recovered from each sample.

### Data analysis and quantification

For HEK293T experiments, fluorescence intensity traces were extracted for each segmented cell and normalized to the fluorescence at the beginning of the recording (*t* = 0) for each individual cell to obtain ΔF/F_0_ values, where F_0_ is the fluorescence intensity at *t* = 0. Single-cell ΔF/F_0_ traces within each imaging dish were averaged to generate a dish-level response trace, and these dish-level traces were used for downstream quantification and statistical analysis. For HyPerRed experiments, fluorescence traces were photobleaching-corrected prior to quantification. Each dish-level trace (representing one biological replicate under a given condition) was fitted to a one-phase exponential decay model using data points acquired during the 0-60 s pre-illumination period, with the fitting curve constrained to start at (0,0). The fitted baseline function was subtracted from the full trace to correct for photobleaching. For ion channel experiments, no photobleaching correction was applied. In miniSOG-GGGS-TRPV1 fusion experiments, a brief illumination-synchronized fluorescence offset^41^ in jRGECO1a coincident with 470 nm light onset and offset was observed and corrected prior to AUC quantification (**Supplementary Figure 9**). This offset was negligible under the acquisition conditions used in other ion channel experiments.

For IMR-90 calcium imaging experiments, ΔF/F_0_ traces were calculated for each individual cell using the normalization procedure described above. Single-cell ΔF/F_0_ traces from each condition were visualized as heatmaps. No photobleaching correction was applied. Per-condition single-cell AUC values were summarized using bar plots, and statistical comparisons were performed using two-tailed *t* tests.

For quantification, the mean ΔF/F_0_ values during 0-60 s were calculated and used as a reference for subsequent quantification. The AUCs between 60 s (illumination onset) and 180 s (or 300 s for experiments with extended stimulation) were calculated relative to this baseline. The peak ΔF/F_0_ was defined as the maximum value within 60-180 s. The onset time was defined as the first time point after 60 s at which ΔF/F_0_ exceeds 0.05 relative to the mean ΔF/F_0_ during 0-60 s. Statistical analyses were performed in GraphPad Prism using two-tailed *t* tests or one-way ANOVA as indicated. For time-course traces, shaded regions represent the standard error of the mean (SEM) across biological replicates. For bar plots, error bars represent the standard deviation (SD).

## Supporting Information

Quantification of HyPerRed and TRP-channel responses, analysis of TRPA1 activation parameters, flow cytometry gating and SYTOX Blue validation for viability analysis, cell viability and cell number measurements under a representative upper-bound HEK293T condition, membrane-targeting and control experiments for miniSOG-induced activation, illumination-synchronized jRGECO1a fluorescence changes, and pharmacological validation of TRPA1-dependent activation in IMR-90 cells.

## Acknowledgments

We thank Ernesto Criado Hidalgo, George Daghlian, Hao Shen, Daniel Tang, and Brian Zhong for helpful discussions; Hao Shen and William Benman for sharing several plasmid constructs; Ann Liu for technical support in cell viability assays; and Akanksha Yadav for technical support in primary cell culture. This research was supported by the National Institutes of Health (DP1 EB033154 to M.G.S.). A.P. was supported by the Swiss National Science Foundation and the Human Frontier Science Program. N.E.I. was supported by the Summer Undergraduate Research Fellowship program at Caltech. M.G.S. is an investigator of the Howard Hughes Medical Institute. We apologize for any unintentional omission of relevant citations.

## Author Contributions

H.L., D.W., and M.G.S. conceived and designed the study. H.L., A.P., N.E.I., and D.W. planned and conducted experiments. H.L. analyzed the data. H.L., D.W., and M.G.S. wrote the manuscript with input from all other authors. M.G.S. supervised the research.

## TOC Graphic

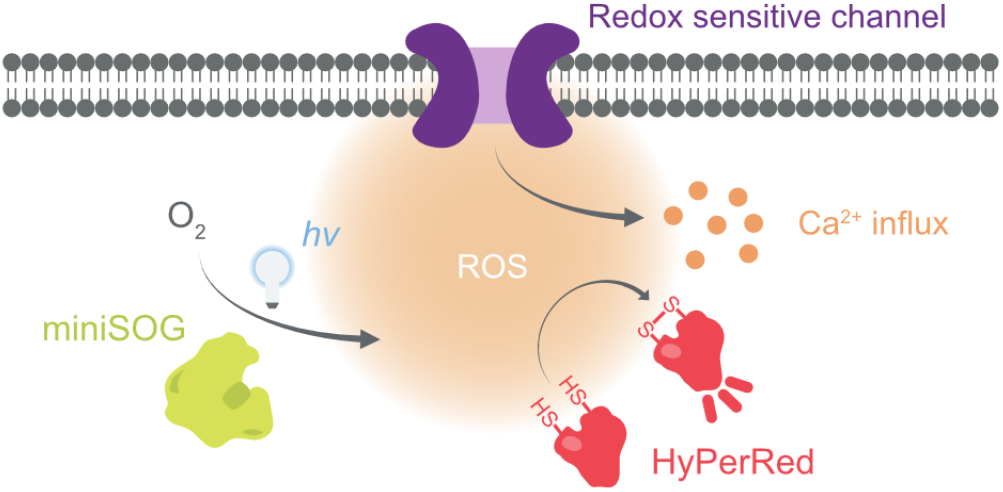

## References

(1) Paulsen, C. E.; Carroll, K. S. Cysteine-Mediated Redox Signaling: Chemistry, Biology, and Tools for Discovery. Chem. Rev. 2013, 113 (7), 4633–4679.

(2) Kaya, A.; Lee, B. C.; Gladyshev, V. N. Regulation of Protein Function by Reversible Methionine Oxidation and the Role of Selenoprotein MsrB1. Antioxid. Redox Signal. 2015, 23 (10), 814–822.

(3) Santos, C. X. C.; Hafstad, A. D.; Beretta, M.; Zhang, M.; Molenaar, C.; Kopec, J.; Fotinou, D.; Murray, T. V.; Cobb, A. M.; Martin, D.; Zeh Silva, M.; Anilkumar, N.; Schröder, K.; Shanahan, C. M.; Brewer, A. C.; Brandes, R. P.; Blanc, E.; Parsons, M.; Belousov, V.; Cammack, R.; Hider, R. C.; Steiner, R. A.; Shah, A. M. Targeted Redox Inhibition of Protein Phosphatase 1 by Nox4 Regulates eIF2α-mediated Stress Signaling. EMBO J. 2016, 35 (3), 319–334.

(4) Poli, G.; Leonarduzzi, G.; Biasi, F.; Chiarpotto, E. Oxidative Stress and Cell Signalling. Curr. Med. Chem. 2004, 11 (9), 1163–1182.

(5) Shu, X.; Lev-Ram, V.; Deerinck, T. J.; Qi, Y.; Ramko, E. B.; Davidson, M. W.; Jin, Y.; Ellisman, M. H.; Tsien, R. Y. A Genetically Encoded Tag for Correlated Light and Electron Microscopy of Intact Cells, Tissues, and Organisms. PLoS Biol. 2011, 9 (4), e1001041.

(6) Bulina, M. E.; Chudakov, D. M.; Britanova, O. V.; Yanushevich, Y. G.; Staroverov, D. B.; Chepurnykh, T. V.; Merzlyak, E. M.; Shkrob, M. A.; Lukyanov, S.; Lukyanov, K. A. A Genetically Encoded Photosensitizer. Nat. Biotechnol. 2005, 24 (1), 95–99.

(7) Makhijani, K.; To, T.-L.; Ruiz-González, R.; Lafaye, C.; Royant, A.; Shu, X. Precision Optogenetic Tool for Selective Single- and Multiple-Cell Ablation in a Live Animal Model System. Cell Chem. Biol. 2017, 24 (1), 110–119.

(8) Takemoto, K.; Matsuda, T.; Sakai, N.; Fu, D.; Noda, M.; Uchiyama, S.; Kotera, I.; Arai, Y.; Horiuchi, M.; Fukui, K.; Ayabe, T.; Inagaki, F.; Suzuki, H.; Nagai, T. SuperNova, a Monomeric Photosensitizing Fluorescent Protein for Chromophore-Assisted Light Inactivation. Sci. Rep. 2013, 3 (1), 2629.

(9) Lin, J. Y.; Sann, S. B.; Zhou, K.; Nabavi, S.; Proulx, C. D.; Malinow, R.; Jin, Y.; Tsien, R. Y. Optogenetic Inhibition of Synaptic Release with Chromophore-Assisted Light Inactivation (CALI). Neuron 2013, 79 (2), 241–253.

(10) Xiong, Y.; Tian, X.; Ai, H.-W. Molecular Tools to Generate Reactive Oxygen Species in Biological Systems. Bioconjug. Chem. 2019, 30 (5), 1297–1303.

(11) Hoerndli, F. J.; Wang, R.; Mellem, J. E.; Kallarackal, A.; Brockie, P. J.; Thacker, C.; Madsen, D. M.; Maricq, V. Neuronal Activity and CaMKII Regulate Kinesin-Mediated Transport of Synaptic AMPARs. Neuron 2015, 86 (2), 457–474.

(12) Grabenbauer, M.; Geerts, W. J. C.; Fernadez-Rodriguez, J.; Hoenger, A.; Koster, A. J.; Nilsson, T. Correlative Microscopy and Electron Tomography of GFP through Photooxidation. Nat. Methods 2005, 2 (11), 857–862.

(13) Ou, H. D.; Kwiatkowski, W.; Deerinck, T. J.; Noske, A.; Blain, K. Y.; Land, H. S.; Soria, C.; Powers, C. J.; May, A. P.; Shu, X.; Tsien, R. Y.; Fitzpatrick, J. A. J.; Long, J. A.; Ellisman, M. H.; Choe, S.; O’Shea, C. C. A Structural Basis for the Assembly and Functions of a Viral Polymer That Inactivates Multiple Tumor Suppressors. Cell 2012, 151 (2), 304–319.

(14) Horstmann, H.; Vasileva, M.; Kuner, T. Photooxidation-Guided Ultrastructural Identification and Analysis of Cells in Neuronal Tissue Labeled with Green Fluorescent Protein. PLoS One 2013, 8 (5), e64764.

(15) Boassa, D.; Lemieux, S. P.; Lev-Ram, V.; Hu, J.; Xiong, Q.; Phan, S.; Mackey, M.; Ramachandra, R.; Peace, R. E.; Adams, S. R.; Ellisman, M. H.; Ngo, J. T. Split-miniSOG for Spatially Detecting Intracellular Protein-Protein Interactions by Correlated Light and Electron Microscopy. Cell Chem. Biol. 2019, 26 (10), 1407–1416.e5.

(16) Kim, D.-I.; Park, S.; Park, S.; Ye, M.; Chen, J. Y.; Kang, S. J.; Jhang, J.; Hunker, A. C.; Zweifel, L. S.; Caron, K. M.; Vaughan, J. M.; Saghatelian, A.; Palmiter, R. D.; Han, S. Presynaptic Sensor and Silencer of Peptidergic Transmission Reveal Neuropeptides as Primary Transmitters in Pontine Fear Circuit. Cell 2024, 187 (18), 5102–5117.e16.

(17) Boassa, D.; Berlanga, M. L.; Yang, M. A.; Terada, M.; Hu, J.; Bushong, E. A.; Hwang, M.; Masliah, E.; George, J. M.; Ellisman, M. H. Mapping the Subcellular Distribution of α-Synuclein in Neurons Using Genetically Encoded Probes for Correlated Light and Electron Microscopy: Implications for Parkinson’s Disease Pathogenesis. J. Neurosci. 2013, 33 (6), 2605–2615.

(18) Wang, P.; Tang, W.; Li, Z.; Zou, Z.; Zhou, Y.; Li, R.; Xiong, T.; Wang, J.; Zou, P. Mapping Spatial Transcriptome with Light-Activated Proximity-Dependent RNA Labeling. Nat. Chem. Biol. 2019, 15 (11), 1110–1119.

(19) Ding, T.; Zhu, L.; Fang, Y.; Liu, Y.; Tang, W.; Zou, P. Chromophore-Assisted Proximity Labeling of DNA Reveals Chromosomal Organization in Living Cells. Angew. Chem. Int. Ed. 2020, 59 (51), 22933–22937.

(20) Zhai, Y.; Huang, X.; Zhang, K.; Huang, Y.; Jiang, Y.; Cui, J.; Zhang, Z.; Chiu, C. K. C.; Zhong, W.; Li, G. Spatiotemporal-Resolved Protein Networks Profiling with Photoactivation Dependent Proximity Labeling. Nat. Commun. 2022, 13 (1), 4906.

(21) Zheng, F.; Yu, C.; Zhou, X.; Zou, P. Genetically Encoded Photocatalytic Protein Labeling Enables Spatially-Resolved Profiling of Intracellular Proteome. Nat. Commun. 2023, 14, 2978.

(22) Hananya, N.; Ye, X.; Koren, S.; Muir, T. W. A Genetically Encoded Photoproximity Labeling Approach for Mapping Protein Territories. Proc. Natl. Acad. Sci. U. S. A. 2023, 120 (16), e2219339120.

(23) Ermakova, Y. G.; Bilan, D. S.; Matlashov, M. E.; Mishina, N. M.; Markvicheva, K. N.; Subach, O. M.; Subach, F. V.; Bogeski, I.; Hoth, M.; Enikolopov, G.; Belousov, V. V. Red Fluorescent Genetically Encoded Indicator for Intracellular Hydrogen Peroxide. Nat. Commun. 2014, 5, 5222.

(24) Yoshida, T.; Inoue, R.; Morii, T.; Takahashi, N.; Yamamoto, S.; Hara, Y.; Tominaga, M.; Shimizu, S.; Sato, Y.; Mori, Y. Nitric Oxide Activates TRP Channels by Cysteine S-Nitrosylation. Nat. Chem. Biol. 2006, 2 (11), 596–607.

(25) Susankova, K.; Tousova, K.; Vyklicky, L.; Teisinger, J.; Vlachova, V. Reducing and Oxidizing Agents Sensitize Heat-Activated Vanilloid Receptor (TRPV1) Current. Mol. Pharmacol. 2006, 70 (1), 383–394.

(26) Wang, S.; Chuang, H.-H. C-Terminal Dimerization Activates the Nociceptive Transduction Channel Transient Receptor Potential Vanilloid 1. J. Biol. Chem. 2011, 286 (47), 40601–40607.

(27) Hernández-Morales, M.; Shang, T.; Chen, J.; Han, V.; Liu, C. Lipid Oxidation Induced by RF Waves and Mediated by Ferritin Iron Causes Activation of Ferritin-Tagged Ion Channels. Cell Rep. 2020, 30 (10), 3250–3260.e7.

(28) Brier, M. I.; Mundell, J. W.; Yu, X.; Su, L.; Holmann, A.; Squeri, J.; Zhang, B.; Stanley, S. A.; Friedman, J. M.; Dordick, J. S. Uncovering a Possible Role of Reactive Oxygen Species in Magnetogenetics. Sci. Rep. 2020, 10 (1), 13096.

(29) Sawada, Y.; Hosokawa, H.; Matsumura, K.; Kobayashi, S. Activation of Transient Receptor Potential Ankyrin 1 by Hydrogen Peroxide. Eur. J. Neurosci. 2008, 27 (5), 1131–1142.

(30) Miyake, T.; Nakamura, S.; Zhao, M.; So, K.; Inoue, K.; Numata, T.; Takahashi, N.; Shirakawa, H.; Mori, Y.; Nakagawa, T.; Kaneko, S. Cold Sensitivity of TRPA1 Is Unveiled by the Prolyl Hydroxylation Blockade-Induced Sensitization to ROS. Nat. Commun. 2016, 7, 12840.

(31) Chuang, H.-H.; Lin, S. Oxidative Challenges Sensitize the Capsaicin Receptor by Covalent Cysteine Modification. Proc. Natl. Acad. Sci. U. S. A. 2009, 106 (47), 20097–20102.

(32) Dana, H.; Mohar, B.; Sun, Y.; Narayan, S.; Gordus, A.; Hasseman, J. P.; Tsegaye, G.; Holt, G. T.; Hu, A.; Walpita, D.; Patel, R.; Macklin, J. J.; Bargmann, C. I.; Ahrens, M. B.; Schreiter, E. R.; Jayaraman, V.; Looger, L. L.; Svoboda, K.; Kim, D. S. Sensitive Red Protein Calcium Indicators for Imaging Neural Activity. eLife 2016, 5, e12727.

(33) Ruiz-González, R.; Cortajarena, A. L.; Mejias, S. H.; Agut, M.; Nonell, S.; Flors, C. Singlet Oxygen Generation by the Genetically Encoded Tag miniSOG. J. Am. Chem. Soc. 2013, 135 (26), 9564–9567.

(34) Wilkinson, F.; Helman, W. P.; Ross, A. B. Rate Constants for the Decay and Reactions of the Lowest Electronically Excited Singlet State of Molecular Oxygen in Solution. An Expanded and Revised Compilation. J. Phys. Chem. Ref. Data 1995, 24 (2), 663–677.

(35) Stokes, A.; Wakano, C.; Koblan-Huberson, M.; Adra, C. N.; Fleig, A.; Turner, H. TRPA1 Is a Substrate for de-Ubiquitination by the Tumor Suppressor CYLD. Cell. Signal. 2006, 18 (10), 1584–1594.

(36) Tonello, R.; Fusi, C.; Materazzi, S.; Marone, I. M.; De Logu, F.; Benemei, S.; Gonçalves, M. C.; Coppi, E.; Castro-Junior, C. J.; Gomez, M. V.; Geppetti, P.; Ferreira, J.; Nassini, R. The Peptide Phα1β, from Spider Venom, Acts as a TRPA1 Channel Antagonist with Antinociceptive Effects in Mice. Br. J. Pharmacol. 2017, 174 (1), 57–69.

(37) Deering-Rice, C. E.; Memon, T.; Lu, Z.; Romero, E. G.; Cox, J.; Taylor-Clark, T.; Veranth, J. M.; Reilly, C. Differential Activation of TRPA1 by Diesel Exhaust Particles: Relationships between Chemical Composition, Potency, and Lung Toxicity. Chem. Res. Toxicol. 2019, 32 (6), 1040–1050.

(38) Xu, C.; He, S.; Wei, X.; Huang, J.; Xu, M.; Pu, K. Activatable Sonoafterglow Nanoprobes for T-Cell Imaging. Adv. Mater. 2023, 35 (30), 2211651.

(39) Wei, X.; Xu, C.; Cheng, P.; Hu, Y.; Liu, J.; Xu, M.; Huang, J.; Zhang, Y.; Pu, K. Leveraging Long-Distance Singlet-Oxygen Transfer for Bienzyme-Locked Afterglow Imaging of Intratumoral Granule Enzymes. J. Am. Chem. Soc. 2024, 146 (25), 17393–17403.

(40) Stringer, C.; Wang, T.; Michaelos, M.; Pachitariu, M. Cellpose: A Generalist Algorithm for Cellular Segmentation. Nat. Methods 2021, 18 (1), 100–106.

(41) Xiang, K. M.; Lampson, H.; Hayward, R. F.; York, A. G.; Ingaramo, M.; Cohen, A. E. Mechanism of Giant Magnetic Field Effect in a Red Fluorescent Protein. J. Am. Chem. Soc. 2025, 147 (21), 18088–18099.

