## Supplementary Information for "Modulating Protein Function through Genetically Encoded Oxidative Chemistry"

Contents:
Supplementary Figure 1-10

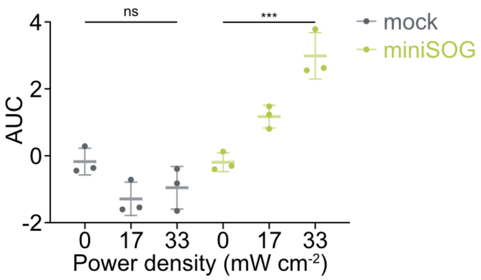


**Supplementary Figure 1. Light-intensity dependence of HyPerRed modulation by miniSOG.**
Quantification of HyPerRed fluorescence responses under different illumination intensities in mock control cells (HyPerRed + empty vector) and miniSOG-expressing HEK293T cells. AUC values were calculated from the time-course traces shown in Figure 1C-E. Fluorescence responses in miniSOG-expressing cells increased with power density, whereas those in control cells did not, indicating that genetically encoded ROS generation depends on illumination intensity. Data are mean ± s.d. (n = 3 independent imaging dishes). Statistical significance was determined by one-way ANOVA with multiple comparisons; ns, not significant; ***, $p < 0.001$.

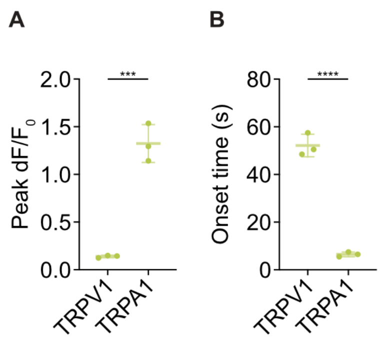


**Supplementary Figure 2. Quantification of TRPA1 and TRPV1 activation induced by miniSOG-derived ROS.**
**(A)** Peak $\Delta F/F_0$ values quantified from the responses in Figure 2 show significantly larger $Ca^{2+}$ amplitudes in miniSOG + TRPA1 cells than in miniSOG + TRPV1 cells.
**(B)** Corresponding activation onset times indicate faster activation kinetics for TRPA1. Data are mean ± s.d. (n = 3 independent imaging dishes). Statistical significance was determined by two-tailed *t* tests; ***, $p < 0.001$; ****, $p < 0.0001$.

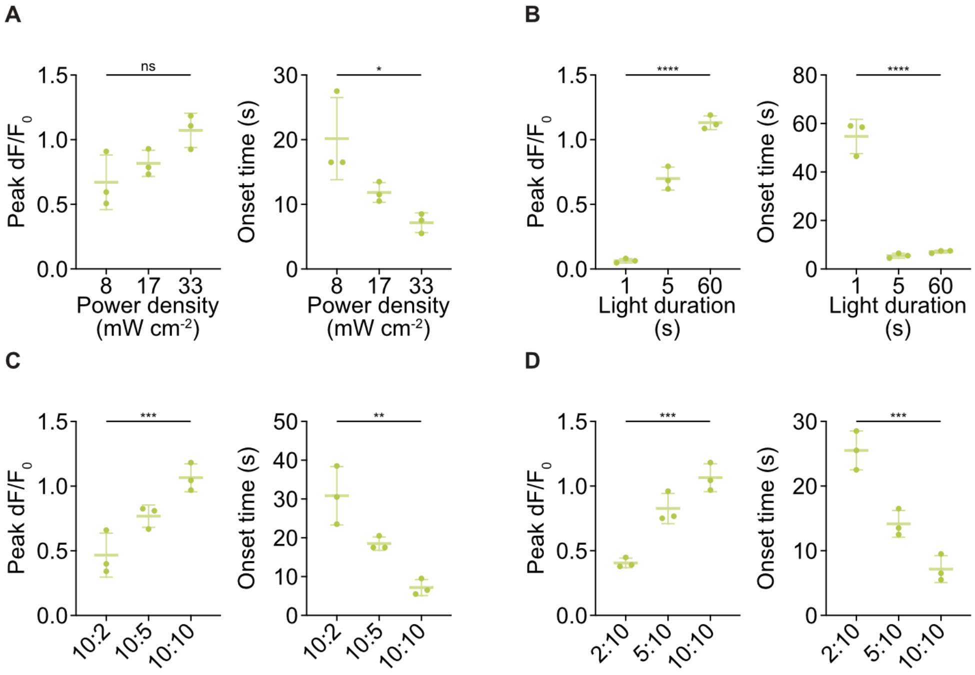


**Supplementary Figure 3. Quantitative analysis of TRPA1 activation parameters under varying optical and genetic conditions.**

**(A)** Quantification of peak $\Delta F/F_0$ and activation onset time under different light intensities corresponding to Figure 3A. While peak amplitudes showed no significant difference, onset time decreased at higher intensities, indicating faster activation kinetics.

**(B)** Peak $\Delta F/F_0$ and onset time under different light durations corresponding to Figure 3B. Both response amplitude increased and onset times decreased with longer illumination.

**(C)** Peak $\Delta F/F_0$ and onset time for cells transfected with fixed TRPA1 and increasing miniSOG plasmid mass, corresponding to Figure 3C. Increasing miniSOG plasmid mass enhanced both response magnitude and activation kinetics.

**(D)** Peak $\Delta F/F_0$ and onset time for cells transfected with fixed miniSOG and increasing TRPA1 plasmid mass, corresponding to Figure 3D. Increasing TRPA1 plasmid mass likewise enhanced response magnitude and activation kinetics. Data are mean ± s.d. (n = 3 independent imaging dishes). Statistical significance was determined by one-way ANOVA with multiple comparisons; ns, not significant; **, $p < 0.01$; ***, $p < 0.001$; ****, $p < 0.0001$.

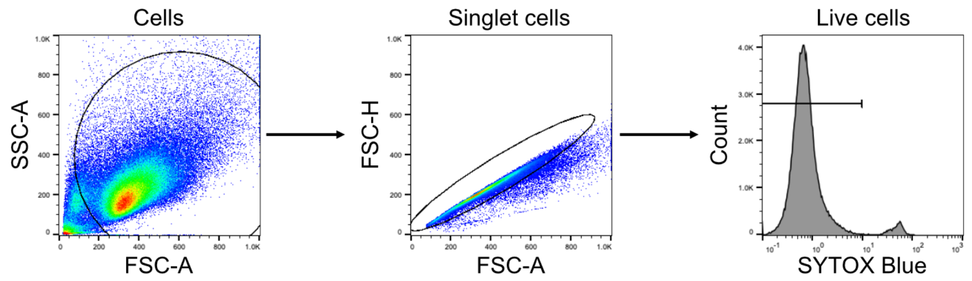


**Supplementary Figure 4. Flow cytometry gating strategy for viability analysis.**
Representative flow cytometry gating strategy used for quantification of cell viability in HEK293T cells expressing miniSOG. Events were first gated on the main cell population using FSC-A and SSC-A, followed by singlet selection using FSC-H versus FSC-A. Live cells were then defined as SYTOX Blue-negative events. The same gating strategy was applied to all samples included in the viability analysis.

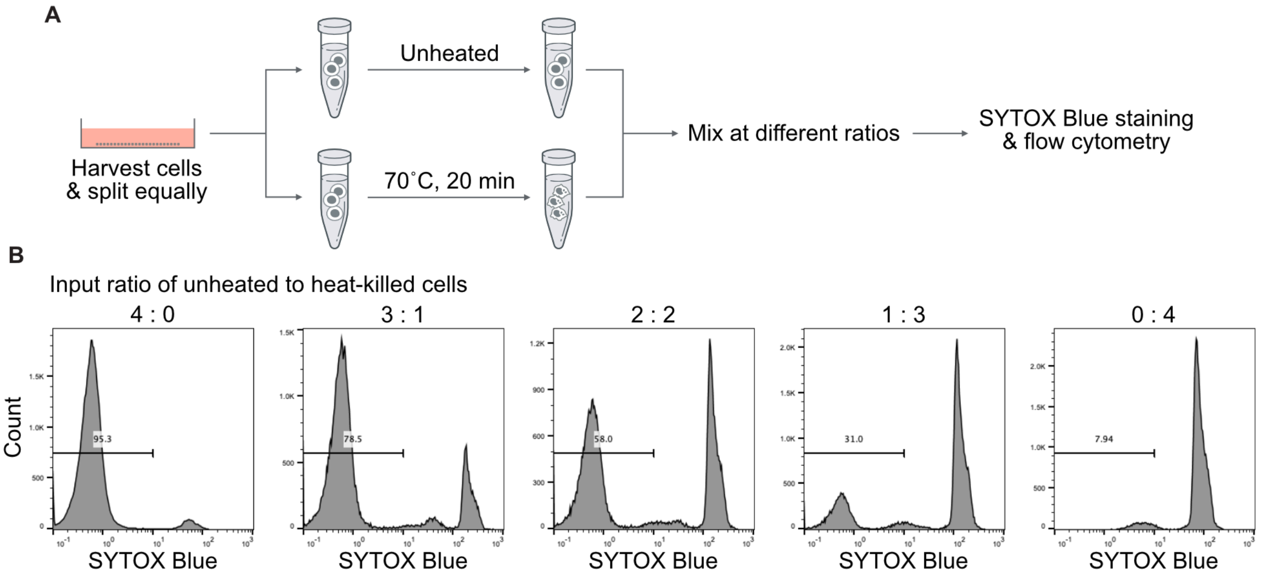


**Supplementary Figure 5. Validation of SYTOX Blue staining for flow-cytometric viability analysis.**
**(A)** Schematic of the preparation of unheated and heat-killed HEK293T cells. Cells were harvested and split equally. One fraction was left unheated, and the other was heat-killed at 70˚C for 20 min. The two fractions were then mixed at the indicated ratios, stained with SYTOX Blue, and analyzed by flow cytometry.
**(B)** Representative SYTOX Blue histograms for mixtures containing increasing fractions of heat-killed cells. The horizontal gate indicates the SYTOX Blue-negative population, and numbers indicate the percentage of events within this gate.

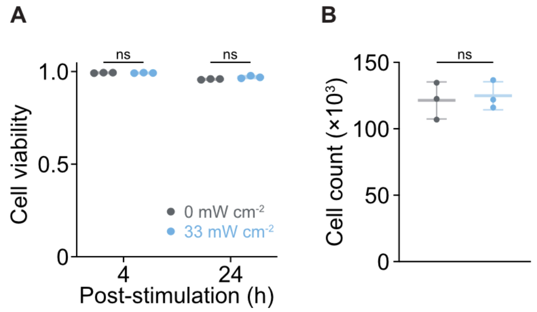


**Supplementary Figure 6. miniSOG expression and blue-light stimulation do not reduce cell viability and cell number under a representative upper-bound HEK293T condition.**

**(A)** Cell viability quantified by flow cytometry using SYTOX Blue in HEK293T cells expressing miniSOG alone at the highest miniSOG DNA input tested in Figure 4C. Cells were analyzed at 4 h and 24 h after blue-light stimulation (33 mW $cm^{-2}$, 60 s) or under 0 mW $cm^{-2}$ control conditions. No significant difference in viability was observed at either time point.

**(B)** Cell number measured at 24 h under the same conditions. No significant difference in cell number was observed between illuminated and non-illuminated cells. Data are mean ± s.d. (n = 3 independent sample dishes). Statistical significance was determined by two-tailed *t* tests; ns, not significant.

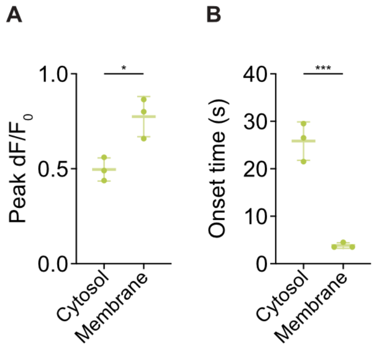


**Supplementary Figure 7. Membrane localization of miniSOG enhances the amplitude and kinetics of TRPA1 activation.**

**(A)** Comparison of the maximum fluorescence change (peak $\Delta F/F_0$) in TRPA1-expressing cells co-expressing cytosolic or membrane-targeted miniSOG under identical illumination (33 mW $cm^{-2}$, 60-120 s).

**(B)** Comparison of activation kinetics (Onset time) under the same conditions. Membrane-targeted miniSOG resulted in significantly shorter onset times. Data are mean ± s.d. (n = 3 independent imaging dishes). Statistical significance was determined by two-tailed *t* tests; *, $p < 0.05$; ***, $p < 0.001$.

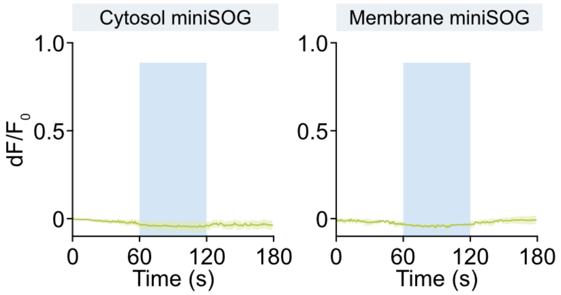


**Supplementary Figure 8. miniSOG alone does not produce detectable $Ca^{2+}$ signals in the absence of TRPA1.** Time-course traces of jRGECO1a fluorescence in HEK293T cells expressing cytosolic (left) or membrane-targeted (right) miniSOG without TRPA1 co-expression. Blue shading indicates the illumination period (33 mW $cm^{-2}$, 60-120 s). No measurable $\Delta F/F_0$ changes were observed under either condition, confirming that $Ca^{2+}$ responses require TRPA1 co-expression. Data are mean ± s. d. (n = 3 independent imaging dishes).

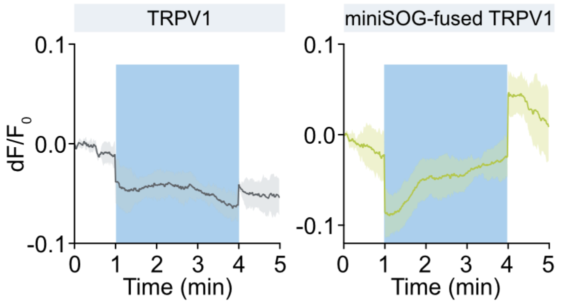


**Supplementary Figure 9. Illumination-synchronized fluorescence changes in jRGECO1a during blue-light stimulation.**

Representative $\Delta F/F_0$ traces of HEK293T cells co-expressing jRGECO1a with TRPV1 (left) or the miniSOG-GGGS-TRPV1 fusion (right) under identical 470 nm illumination (66 mW $cm^{-2}$, 1-4 min). A brief fluorescence decrease at illumination onset and a corresponding increase at illumination offset were observed in both conditions, synchronized with the onset and removal of blue light. Such illumination-dependent fluorescence changes are consistent with previously reported light-induced dark-state behavior of red fluorescent proteins[1]. To account for this effect, traces used for AUC quantification in Figure 6 were corrected for the illumination-synchronized offset. The difference between the fusion construct and control responses remains unchanged after correction.

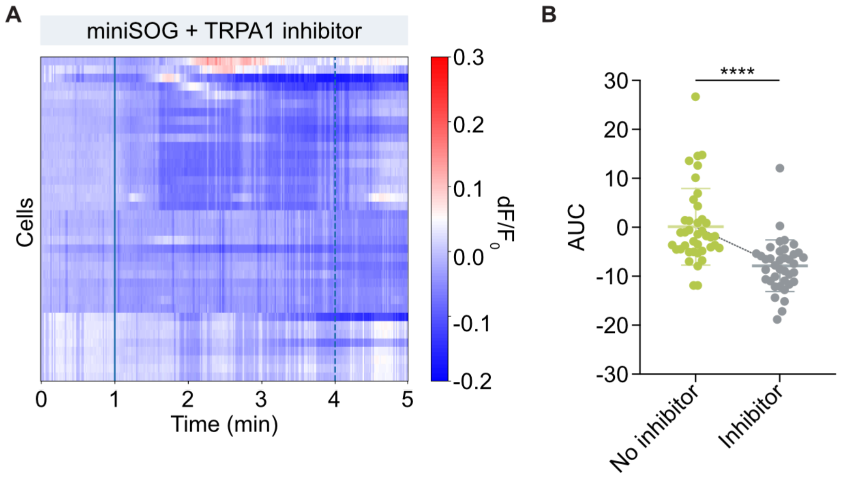


**Supplementary Figure 10. Pharmacological inhibition confirms TRPA1-dependent activation induced by miniSOG-generated ROS.**

**(A)** Single-cell $Ca^{2+}$ response heatmaps for miniSOG-expressing IMR-90 cells treated with a TRPA1 inhibitor HC-030031 (10 μM) during blue light illumination (66 mW $cm^{-2}$, 1-4 min). Treatment with the TRPA1 inhibitor markedly suppressed the $Ca^{2+}$ responses observed in miniSOG-expressing cells.

**(B)** Quantification of $Ca^{2+}$ responses shown as integrated fluorescence (AUC) in the presence or absence of TRPA1 inhibitor. Data are mean ± s.d. (no inhibitor: n = 40 cells; inhibitor: n = 38 cells). Statistical significance was determined by two-tailed *t* tests; ****, $p < 0.0001$.

### References


(1) Xiang, K. M.; Lampson, H.; Hayward, R. F.; York, A. G.; Ingaramo, M.; Cohen, A. E. Mechanism of Giant Magnetic Field Effect in a Red Fluorescent Protein. *J. Am. Chem. Soc.* **2025**, *147* (21), 18088–18099.
